# Endoplasmic reticulum phospholipid scramblase activity revealed after protein reconstitution into giant unilamellar vesicles containing a photostable lipid reporter

**DOI:** 10.1101/2021.04.01.437912

**Authors:** Patricia P. M. Mathiassen, Anant K. Menon, Thomas Günther Pomorski

**Author notes:** **Author for correspondence** Anant K. Menon, Email address and Thomas Günther Pomorski.

## Abstract

Transbilayer movement of phospholipids in biological membranes is mediated by a diverse set of lipid transporters. Among them are scramblases that facilitate a rapid bi-directional movement of lipids without metabolic energy input. Here, we established a new fluorescence microscopy-based assay for detecting phospholipid scramblase activity of membrane proteins upon their reconstitution into giant unilamellar vesicles formed from proteoliposomes by electroformation. The assay is based on chemical bleaching of fluorescence of a photostable ATTO-dye labeled phospholipid with the membrane-impermeant reductant sodium dithionite. We demonstrate that this new methodology is suitable for the study of the scramblase activity of the yeast endoplasmic reticulum at single vesicle level.

## Introduction

The lipid distribution across cellular membranes is regulated by a diverse set of membrane transporters that control the movement of lipids across membranes. These transporters can be classified into two categories: (i) ATP-driven, vectorial transporters that actively translocate lipids from one membrane leaflet to the other, often with high specificity, and (ii) ATP-independent transporters, also called scramblases that facilitate a rapid bi-directional movement of lipids without metabolic energy input. Scramblases are either constitutively active or regulated by physiological stimuli, e.g. a rise in intracellular Ca^2+^ or proteolytic cleavage ^1,2^. Constitutively active scramblases are found in the bacterial cytoplasmic membrane and the endoplasmic reticulum (ER) and promote uniform growth of the membranes after synthesis of lipids on the cytoplasmic side. The molecular identity of specific ER scramblases is not known, but constitutive phospholipid scramblase activity is an unexpected property of Class A G protein-coupled receptors ^3–5^. In the plasma membrane of eukaryotic cells, regulated scramblases are responsible for a controlled loss of lipid asymmetry and the appearance of the anionic phospholipid phosphatidylserine at the cell surface. Two major families of regulated scramblases have been identified: the TMEM16 family ^6^ and the Xk-related (Xkr) family ^7^.

The mechanistic analysis of lipid scramblases at the molecular level is challenging due to the complexity of the membrane in which they are embedded. Current analyses are based on their reconstitution into large unilamellar liposomes (LUVs, 50–500 nm diameter) of desired bulk lipid composition followed by ensemble averaged biophysical or biochemical studies. These studies provided the first insights into the characteristics of scramblase activities. For example, the ER scrambling activity displays a relatively low specificity and transports glycerophospholipids as well as ceramide-based lipids equally well within the limited time-resolution of the activity assays ^8,9^. The reconstitution studies show unambiguously that not all proteins can scramble lipids ^10,11^ and protein modification studies suggest that there are at least two proteins that contribute to overall scramblase activity in the ER ^12^.

A significant drawback of ensemble measurements arises from the compositional heterogeneity of proteoliposome reconstitutions, which hampers quantitative analysis of vesicle properties (including stoichiometry of lipids, sterols, transmembrane proteins) and correlation with protein activity ^13^. Giant unilameller vesicles (GUVs, 10-100 µm diameter) are model membrane systems with dimensions comparable to that of a cell, allowing single vesicle analysis directly by microscopy techniques such as fluorescence microscopy, fluorescence correlation spectroscopy or atomic force microscopy ^14–17^. Furthermore, such vesicles can be micro-manipulated for position control or mechanical probing. GUVs have been extensively used in membrane biophysics ^18,19^, and in studies of cargo inclusion into the GUV lumen ^20,21^, and incorporation of transporter proteins into the membrane ^22,23^. Although several protocols have proven successful for studying various membrane transporters, only one assay has been established for studying scramblase activity in GUVs. This assay is based on detecting scramblase activity by observing vesicle shape changes upon addition of external lipids ^24^. However, the approach requires tight osmotic control of buffers and stable surroundings, while additionally resulting in a low quantitative output. In this study, we established a new fluorescence microscopy-based assay for detecting phospholipid scramblase activity. The protocol allows generation and imaging of individual protein-containing GUVs and was employed to visualize the scramblase activity of ER membrane proteins. We performed reconstitution under low salt conditions that promote mild electroformation of GUVs from preformed proteoliposomes ^22,25^. Our work sets the stage for future experiments where the critical question of the effect of membrane lipid composition on scramblase activity can be examined by generating GUVs using a combination of proteoliposomes reconstituted with purified scramblases such as members of the GPCR, TMEM16 and Xkr8 families, and LUVs of the desired composition ^26,27^.

## Results

### ATTO488-phosphatidylethanolamine (ATTO488-PE) is a photostable reporter of phospholipid scramblase activity

Fluorescence-based phospholipid scramblase assays are typically performed using an ensemble of LUVs containing reporter lipids modified with the nitrobenzoxadiazole (NBD) fluorophore (Fig. 1a). While effective in cuvette-based applications ^3,12,28,29^, NBD is highly susceptible to photobleaching and therefore impractical for use in the microscope. In contrast, ATTO dyes are photostable (Fig. 1b), making them an obvious choice for fluorescence microscopy applications ^30^. We recently showed that ATTO488 can be rendered non-fluorescent by chemical reduction with the membrane-impermeant dianion dithionite ^31^, suggesting that ATTO488-modified phospholipids could be suitable reporters of scramblase activity in microscopy-based assays. We tested this possibility in a cuvette-based assay as follows: Proteoliposomes were reconstituted using egg phosphatidylcholine, trace amounts of ATTO488-PE and biotinylated phosphatidylethanolamine (biotin-PE), and Triton X-100-solubilized yeast ER membrane proteins (‘Triton Extract’ or TE), as described previously ^9,11^. In these and other experiments, we generally used TE647, a preparation of TE in which the proteins had been fluorescently labeled with an amine-reactive derivative of Alexa Fluor 647 (Suppl. Fig. S1). Protein-free (empty) vesicles were prepared in parallel. We used low salt conditions as these would facilitate the conversion of proteoliposomes to GUVs in larger quantities ^22,25^. Upon adding dithionite to symmetrically labeled vesicles lacking a scramblase, we expected to observe ~50% reduction in fluorescence, as ATTO488-PE molecules located in the outer leaflet of the vesicles are bleached whereas those in the inner leaflet are inaccessible to dithionite (Fig. 1a). However, if the vesicles contain a functional scramblase, ATTO488-PE molecules will be translocated from the inner leaflet to the external leaflet and vice versa, resulting in 100% fluorescence reduction. Our results support this scheme. We found that ATTO488-PE fluorescence was reduced by 55 ± 4.5% (mean ± s.d., 4 replicates) on dithionite treatment of protein-free liposomes (Fig. 1c), consistent with the expectation that the lipids are symmetrically distributed between the two leaflets. In contrast, fluorescence was reduced by 74 ± 4.3% (3 replicates) in TE647-containing proteoliposomes (Fig. 1c). Notably, ATTO488-PE is reduced with a ~5-fold smaller half-time compared to a phospholipid with an NBD-modified acyl chain (Fig. 1d). Despite preparing proteoliposomes at a high protein-to-phospholipid ratio of 22 mg mmol^−1^ where we would expect three scramblases per vesicle on average ^9^, we did not observe the expected 100% reduction in fluorescence. This suggests that not all liposomes are reconstituted with an active scramblase, as also noted in several studies ^32,33^. Notably, previous studies utilizing encapsulated water-soluble 2-NBD-glucose showed that dithionite cannot pass through the membrane of liposomes or proteoliposomes in conditions similar to ours ^4,34^. This indicates that the greater extent of fluorescence reduction seen in proteoliposomes versus liposomes is not due to dithionite permeation into the vesicles. It is rather due to the presence of a scramblase that is able to exchange inner leaflet lipid reporters with the outer leaflet. Based on these observations we conclude that ATTO488-PE is a suitable reporter of scramblase activity in LUVs, with the high photostability needed for microscopy-based assays.

**Figure 1.**
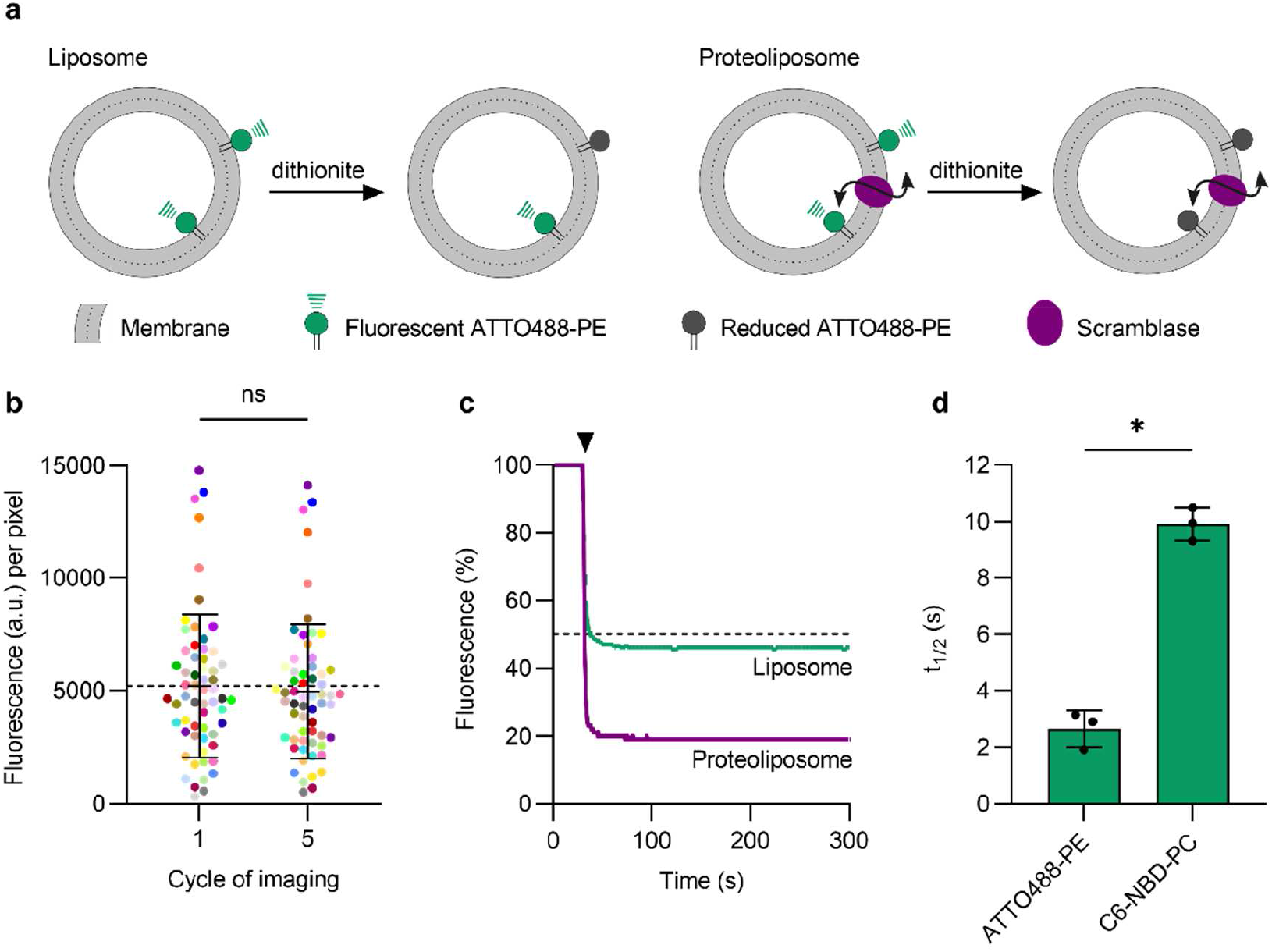
ATTO488-PE is an effective reporter of phospholipid scrambling in large unilamellar vesicles reconstituted with ER membrane proteins. (**a**) Schematic illustration of the scramblase activity assay. The assay makes use of large unilamellar vesicles that contain a trace amount of ATTO488-PE distributed equally between both leaflets and exploits the ability of dithionite to chemically reduce the ATTO488 fluorophore, thereby irreversibly eliminating its fluorescence. In protein-free liposomes (left), ATTO488-PE in the outer leaflet is bleached upon addition of membrane-impermeant dithionite, whereas the inner leaflet pool is protected, resulting in 50% loss of fluorescence. For proteoliposomes with a functional scramblase (right), ATTO488-PE exchanges rapidly between the leaflets, resulting in 100% loss of fluorescence upon dithionite addition. (**b**) Photostability of ATTO488-PE in GUVs. Fluorescence intensity of individual ATTO488-PE labelled GUVs (color-matched between the image cycles, n=60) before and after five confocal imaging scans. The mean ± s.d. is indicated. (**c**) Representative traces showing fluorescence loss on adding dithionite to protein-free liposomes (green) and proteoliposomes (purple) reconstituted with ER membrane proteins. The arrowhead indicates dithionite addition. The horizontal dashed line corresponds to 50% loss of fluorescence. (**d**) Half-time of dithionite-mediated fluorescence reduction of ATTO488-PE and C6-NBD-PC reconstituted into liposomes. Results are presented as mean ± s.d. of at least three independent reconstitutions (unpaired *t* test with Welch’s correction, \**P* < 0.0005).

### Formation of pGUVs containing ATTO488-PE and fluorescent ER membrane proteins

GUVs formed from proteoliposomes by electroformation have been shown to incorporate and maintain scramblase activity as evinced by a shape-change assay ^24^. We therefore used electroformation to generate protein-containing GUVs (pGUVs) from ATTO488-PE-containing proteoliposomes reconstituted with TE647. Briefly, droplets of a proteoliposome suspension, diluted in distilled water, were deposited on two indium tin oxide (ITO) coated glass slides and dehydrated to form a thin lipid film (Fig. 2). A chamber was created by sandwiching the slides, with the lipid-film-bearing surfaces facing each other. The ITO slides were separated with a Teflon spacer with copper electrodes on each side. For electroformation, 250 mM sucrose solution was injected, and the chamber was exposed to an oscillating field resulting in the formation of pGUVs. Empty GUVs (eGUVs) were prepared in parallel from protein-free liposomes. The GUV suspension was removed from the chamber and diluted five-fold in osmotically matched 250 mM glucose solution (Fig. 3a). The procedure promotes the ability of the GUVs to sediment for convenient observation. We obtained a high yield of GUVs, as visualized initially via fluorescence of the ATTO488-PE membrane marker (Fig. 3b, top panels), and subsequently checked for fluorescence of Alexa647, indicative of the presence of reconstituted proteins (Fig. 3b, bottom panels). All GUVs generated from proteoliposomes contained protein (Fig. 3b). The size distribution of the pGUVs and eGUVs was similar (Fig. 3c), with a modal diameter in the range 20-40 µm for both populations. Although the majority of the pGUVs and eGUVs were unilamellar (89 ± 7.6%, n=342 and 66 ± 1.0%, n=340, respectively), we noted occasional GUVs with internal tubular structures, and these were more frequently seen in eGUV preparations (Fig. 3d). GUVs with encapsulated multilamellar or multivesicular vesicles represented only a small fraction of the population (Fig. 3d). These observations are consistent with previous reports showing that application of a thin lipid film on a surface exposed to an electric field results in a high fraction of unilamellar GUVs ^25^. Because of the inaccessible internal bilayers of multilamellar or multivesicular, the latter GUV types were excluded from our analyses.

**Figure 2.**
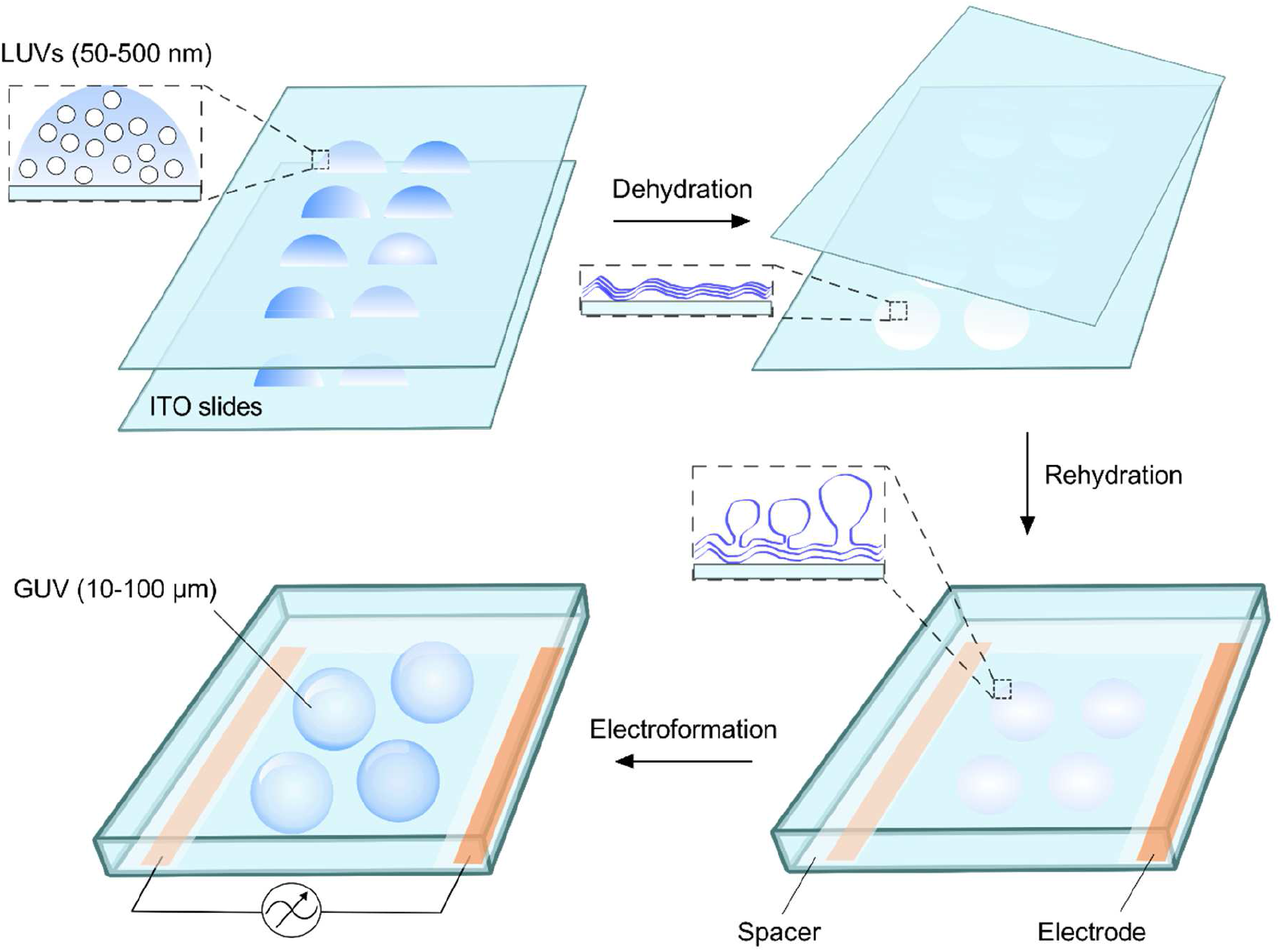
Electroformation of GUVs. Liposomes or proteoliposomes, diluted in distilled water, are applied in droplets on the conductive surface of each of two glass slides coated with indium tin oxide (ITO). The solution is dehydrated overnight in a chamber containing saturated sodium chloride, resulting in the formation of a thin lipid film. The two slides are then assembled into a chamber so that the conducting surfaces with the lipid film face each other, separated by a Teflon spacer containing copper electrodes on each side. The chamber is sealed and held together by clamps (not illustrated), before being connected to a power source and exposed to an oscillating electric field upon injection of 250 mM sucrose solution, resulting in GUV formation.

**Figure 3.**
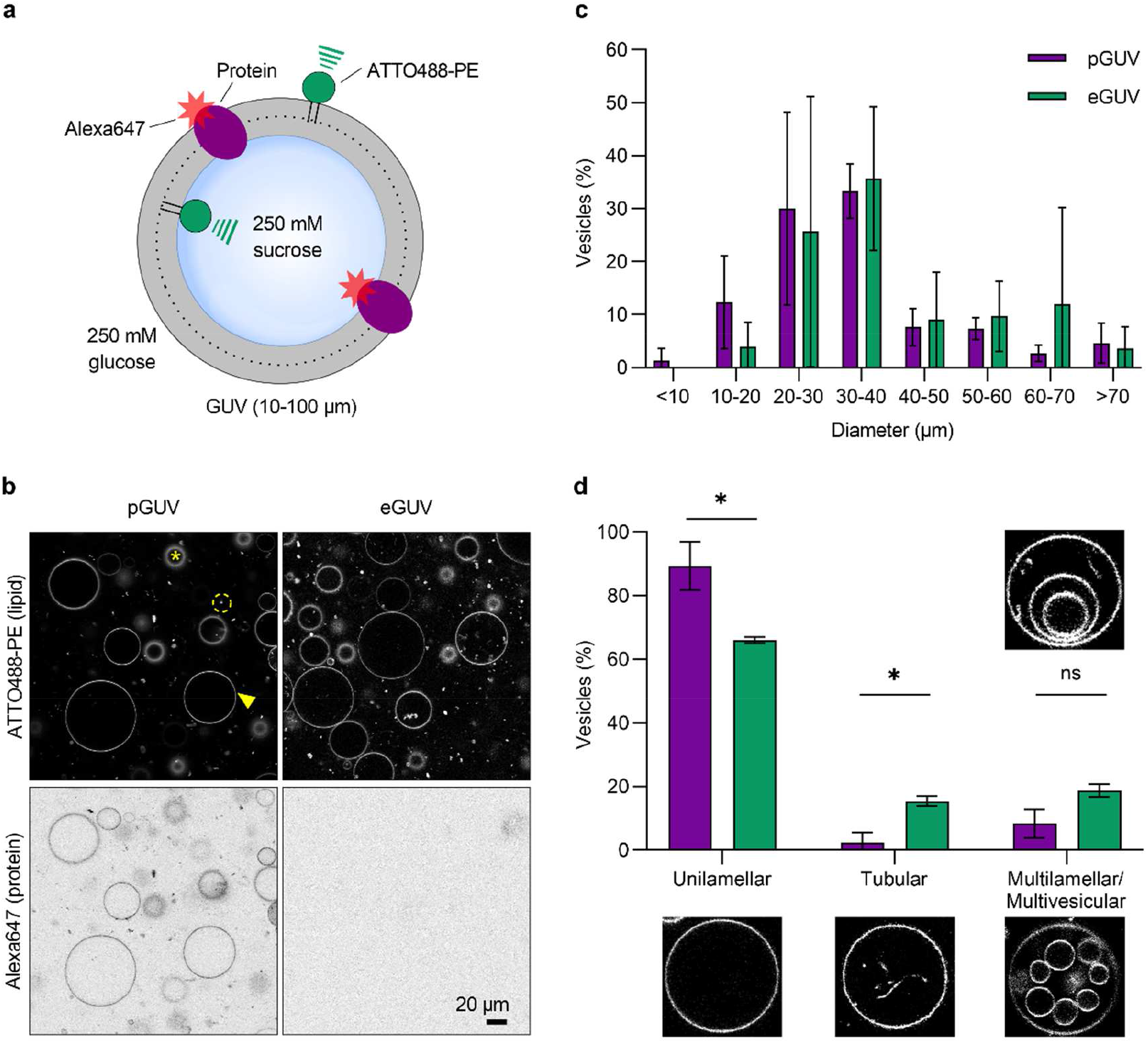
Characteristics of GUVs. (**a**) Protein-containing GUVs (pGUVs) generated in 250 mM sucrose solution were diluted in 250 mM glucose solution for observation. (**b**) Confocal images of pGUVs and empty (protein-free) GUVs (eGUVs) showing fluorescence channels corresponding to ATTO488-PE and Alexa647-labeled ER proteins). The arrowhead indicates an exemplary GUV used for analysis. The asterix indicates an out-of-focus GUV and the dashed circle indicates a free floating proteoliposome. (**c, d**) Size and morphology distribution of pGUVs (purple, n=342) and eGUVs (green, n=340). Data are presented as mean ± s.d. of three technical replicates (two-way ANOVA with Šídák’s multiple-comparisons test, \**P* < 0.05).

### Phospholipid scrambling in GUVs detected by dithionite-mediated bleaching of ATTO488-PE

Next, we sought to investigate scramblase activity in pGUVs. To establish the necessary protocols, we first tested the reactivity of ATTO488-PE to dithionite in eGUVs. A trace amount of biotin-PE was included in preparation of eGUVs to promote their immobilization on avidin-biotin-PEGylated cover glass slides in an incubation chamber (Fig. 4a). The GUVs were imaged via their ATTO488-PE fluorescence (Fig. 4c, t=0). A small volume (1% of the aqueous volume in the chamber) of dithionite solution was added gently to the chamber (Fig. 4a, bottom panel), without overt mixing in order not to disturb the settled/attached GUVs. A time series of fluorescence images was captured over the next 12 min (Fig. 4c, upper panels). Buffer treatment was performed in parallel (Fig. 4c, lower panels). The ATTO488-PE fluorescence intensity of individual eGUVs before, and at different time points after dithionite or buffer addition, was quantified by image analysis using ImageJ as follows (Fig. 4b). An outline of individual GUVs was manually defined based on the fluorescence of ATTO488-PE. Membrane fluorescence was quantified by a circular region of interest (ROI) around each GUV, measuring the integrated density value (Fig. 4b). A ROI in the lumen of the individual GUV was used to quantify the background signal per pixel, which was then subtracted from the membrane signal by multiplying with the area of the membrane ROI. The background signal was typically insignificant but was nevertheless used as an offset correction as part of our standard procedure. We observed some movement of the eGUVs on addition of either buffer or dithionite. The movement occurred within and along the focus plane on the microscope slide indicating that the GUVs were not completely immobilized via the avidin-biotin-PEG system. Nevertheless, it was possible to track a sufficient number of individual GUVs that stayed within the field of view due to sedimentation, as indicated in Fig. 4c, and to quantify their ATTO488 fluorescence before and after dithionite treatment (Fig. 4b, d).

**Figure 4.**
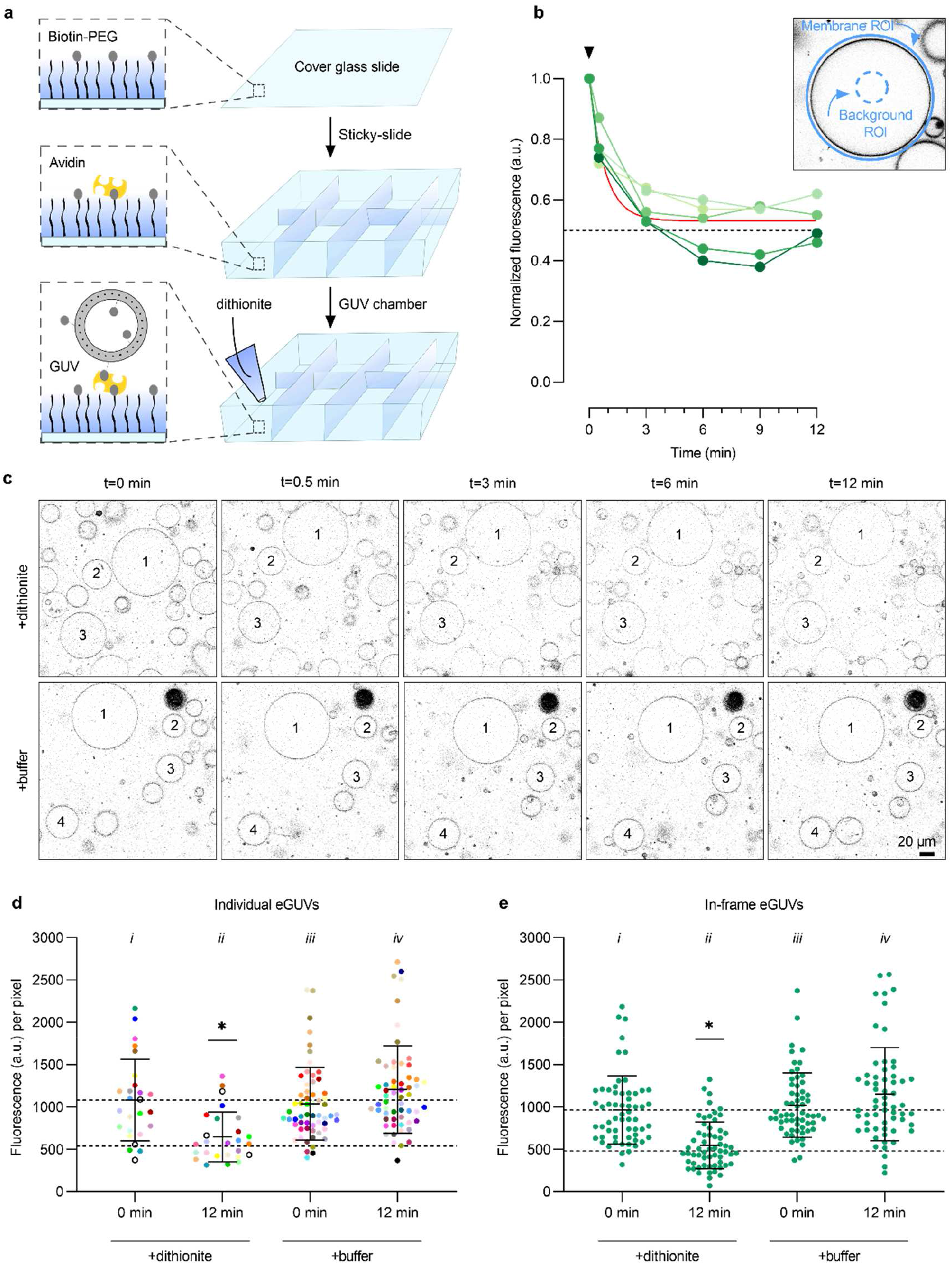
Dithionite bleaching of ATTO488-PE in empty GUVs. (**a**) Microscope chambers were assembled by gluing a sticky-slide onto a biotin-PEGylated cover glass slide to which avidin was added. Unbound avidin was washed away before the GUV suspension was added and observed via confocal microscopy. Dithionite (or buffer) was added to the field of view once the GUVs had sedimented. (**b**) Dithionite reduction of ATTO488-PE fluorescence of five eGUVs (coded in different shades of green). The arrowhead indicates dithionite addition. The red line is a monoexponential fit to the combined data (t_½_=0.56 min, span=0.47). *Inset*, protocol for quantification of ATTO488-PE membrane fluorescence. An outer region of interest (ROI) is placed around the GUV membrane and its total fluorescence is determined. Next, a ROI is defined within the GUV lumen to quantify background fluorescence. The average fluorescence per pixel of the lumenal ROI is determined and scaled to the pixel area of the outer ROI for background subtraction. (**c**) Confocal images of ATTO488-PE-containing eGUVs before (t=0) and at different times after dithionite addition (t=0.5, 3, 6, 12 min). Identical eGUVs (numbered) were followed over the time series. (**d**) Dot plot of ATTO488-PE fluorescence from individually tracked eGUVs (color-matched between the time points) before and 12 min after dithionite (n=26) or buffer treatment (n=60). Black open circles indicate eGUVs that did not react to dithionite. (**e**) Scatter plot of ATTO488-PE fluorescence from all individual eGUVs within the field of view before and 12 min after dithionite and buffer treatment (n=60). Black lines indicate mean ± s.d. (one-way ANOVA with Šídák’s multiple-comparisons test, *\*P* < 0.05). The horizontal dashed lines are provided as a guide to indicate 100% and 50% levels of fluorescence based on the mean value at t=0.

Addition of dithionite caused rapid (t_½_ = 0.58 ± 0.23 min, n=5) reduction of ATTO488-PE fluorescence in individual eGUVs (Fig. 4b), reaching a plateau of 0.53 ± 0.08 (n=5), i.e., a 47% reduction in fluorescence. We extended this exemplary analysis to a larger population of eGUVs. Fig. 4c shows end-point fluorescence (t=12 min) compared with the starting value for individual eGUVs (n=23) in samples treated with either dithionite or buffer; a time course for the dithionite-treated samples is shown in Suppl. Fig. S2. This larger data set (n=23) yielded values comparable to that of our initial analyses of approximately 44% fluorescence reduction, with t_1/2_ = 0.82 min (Fig. 4d, Suppl. Fig. S2). A few eGUVs did not react to dithionite (Fig. 4d and Suppl. Fig. S2, 3 of out 26, black open circles), suggesting uneven diffusion of the reagent over the slide. The ATTO488 fluorescence of eGUVs was not affected when buffer was applied instead of dithionite (Fig. 4d, compare (iv) versus (iii)). As noted above, physical displacement of GUVs on addition of dithionite or buffer preventing convenient tracking of a large number of individual eGUVs. Therefore, in order to increase the sample size for analysis, all eGUVs within the field of view before and after dithionite treatment were analyzed (Fig. 4e). Here, 43% (n=60) of the ATTO488-PE fluorescence was reduced upon dithionite treatment (Fig. 4e, compare (ii) versus (i)), which is comparable to the results of individual tracked vesicles. As expected, buffer treatment did not affect the fluorescence (Fig. 4d, compare (iv) versus (iii)). These results indicate that the extent to which dithionite bleaches ATTO488-PE in eGUVs can be reliably determined not only by tracking individual GUVs, but also by analyzing any in-focus GUVs. The latter approach allows more vesicles to be counted for each replication, thus greatly expanding the GUV sample size if needed.

Before proceeding to investigation of the behavior of ATTO488-PE in pGUVs in response to dithionite, we first established that dithionite does not permeate across the membrane of these vesicles on account of their protein content. To this end, we employed the polar fluorophore Alexa Fluor 647 hydrazide. In solution, this fluorophore reacts readily with dithionite (Fig. 5a), confirming that it is a good reporter of dithionite permeation across the pGUV membrane. pGUVs were generated via electroformation in the presence of Alexa Fluor 647 hydrazide for luminal inclusion and treated with 10 mM dithionite before observation in the microscope. Quantification of the luminal signal 12 min after treatment in comparison with buffer treated pGUVs (Fig. 5b), showed that the luminal marker was protected (Fig. 5c) in all but a minor fraction (<10%) of pGUVs which had somewhat decreased Alexa Fluor 647 hydrazide fluorescence. We conclude that dithionite cannot enter pGUVs on the time scale of our experiments.

**Figure 5.**
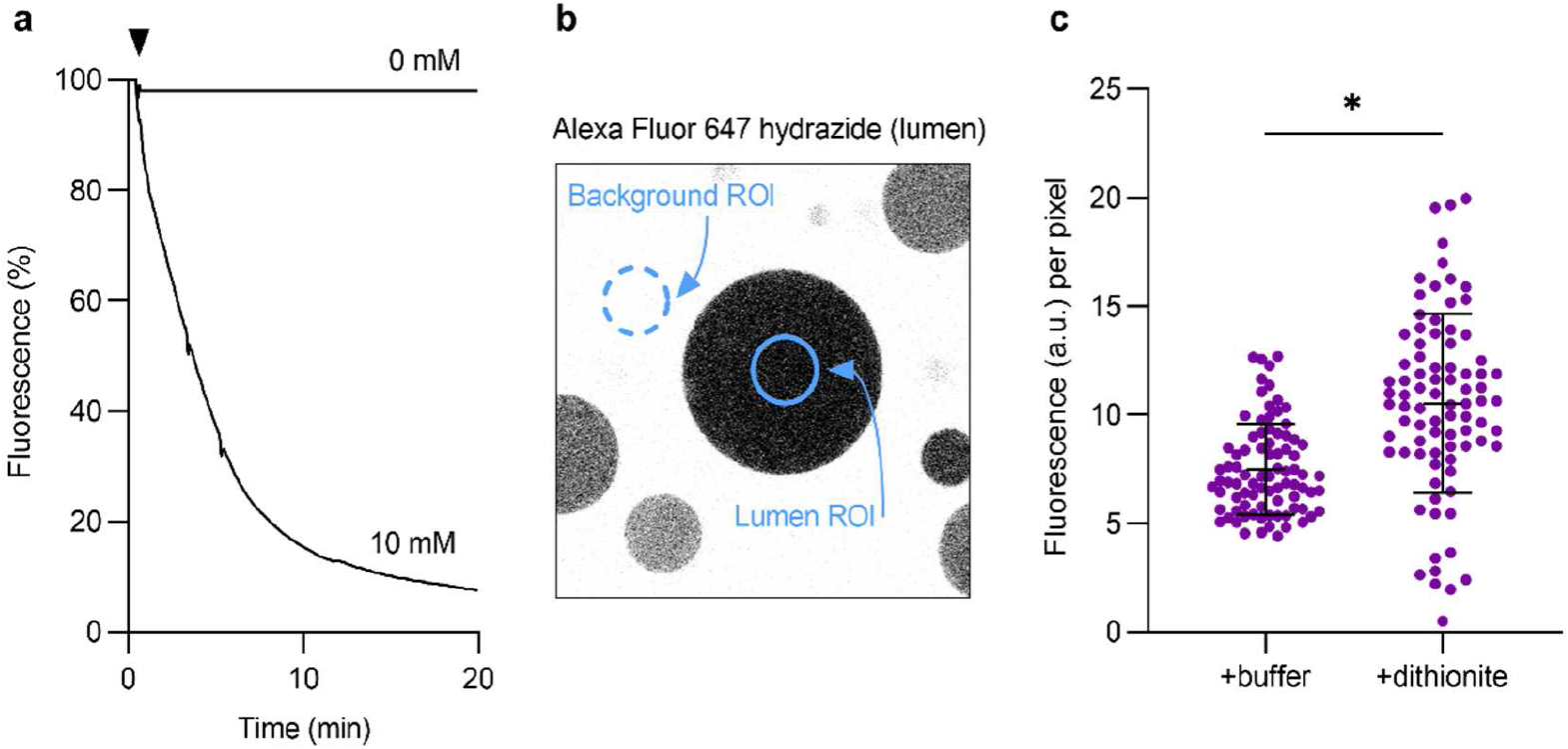
Dithionite does not permeate across the membrane of pGUVs. (**a**) Fluorescence time trace of 10 µM Alexa Fluor 647 hydrazide in 250 mM sucrose solution upon addition of 10 mM dithionite. A buffer control (0 mM dithionite) is shown for comparison. The arrowhead indicates dithionite or buffer addition. (**b**) Confocal image of pGUVs with trapped Alexa Fluor 647 hydrazide. A ROI was placed within the GUV for fluorescence quantification, from which the mean of four background ROIs was subtracted. (**c**) Dot plot of Alexa Fluor 647 hydrazide fluorescence of individual GUVs 12 min after buffer or dithionite addition (n=84). Black lines indicate mean ± s.d. (unpaired *t* test with Welch’s correction, *\*P* < 0.0001).

Our results thus far indicate that dithionite treatment of ATTO488-PE-containing eGUVs produces the expected partial loss of fluorescence, close to 50%, and that dithionite does not permeate across the GUV membrane, irrespective of whether the GUV contains reconstituted proteins. We next went on to treat ATTO488-PE-containing pGUVs with dithionite to assay scrambling of the fluorescent lipid reporter. We expected to produce three populations of pGUVs upon dithionite treatment: (i) a minor population of vesicles that are not affected by dithionite, likely because of uneven spreading of the applied reagent as discussed above, (ii) a population exhibiting ~43-47% lower fluorescence of ATTO488-PE (compared with untreated samples) because of lack of a functional scramblase (as for eGUVs), and (iii) a ‘dark’ population in which all ATTO488-PE is reduced at the end-point of the assay because of the presence of a functional scramblase. The latter vesicles would only be detected as kinetic intermediates exhibiting partial ATTO488-PE fluorescence at early time points.

Fig. 6a shows a time series of fluorescence images of ATTO488-PE-containing pGUVs captured over 12 min after dithionite or buffer addition. Quantification of the percentage of ATTO488-PE-positive GUVs revealed that essentially 100% of eGUVs but only ~29% of pGUVs remained detectable above background levels 12 min after application of dithionite (Fig. 6b). Alexa Fluor 647 hydrazide-loaded pGUVs served as control in these experiments to exclude dislodgement or destruction of pGUVs by dithionite treatment (Fig. 6c). We conclude that whereas the average number of pGUVs in the field of view was the same irrespective of dithionite treatment (Fig. 6c), ~71% of the population had completely lost their ATTO488-PE fluorescence by t=12 min.

**Figure 6.**
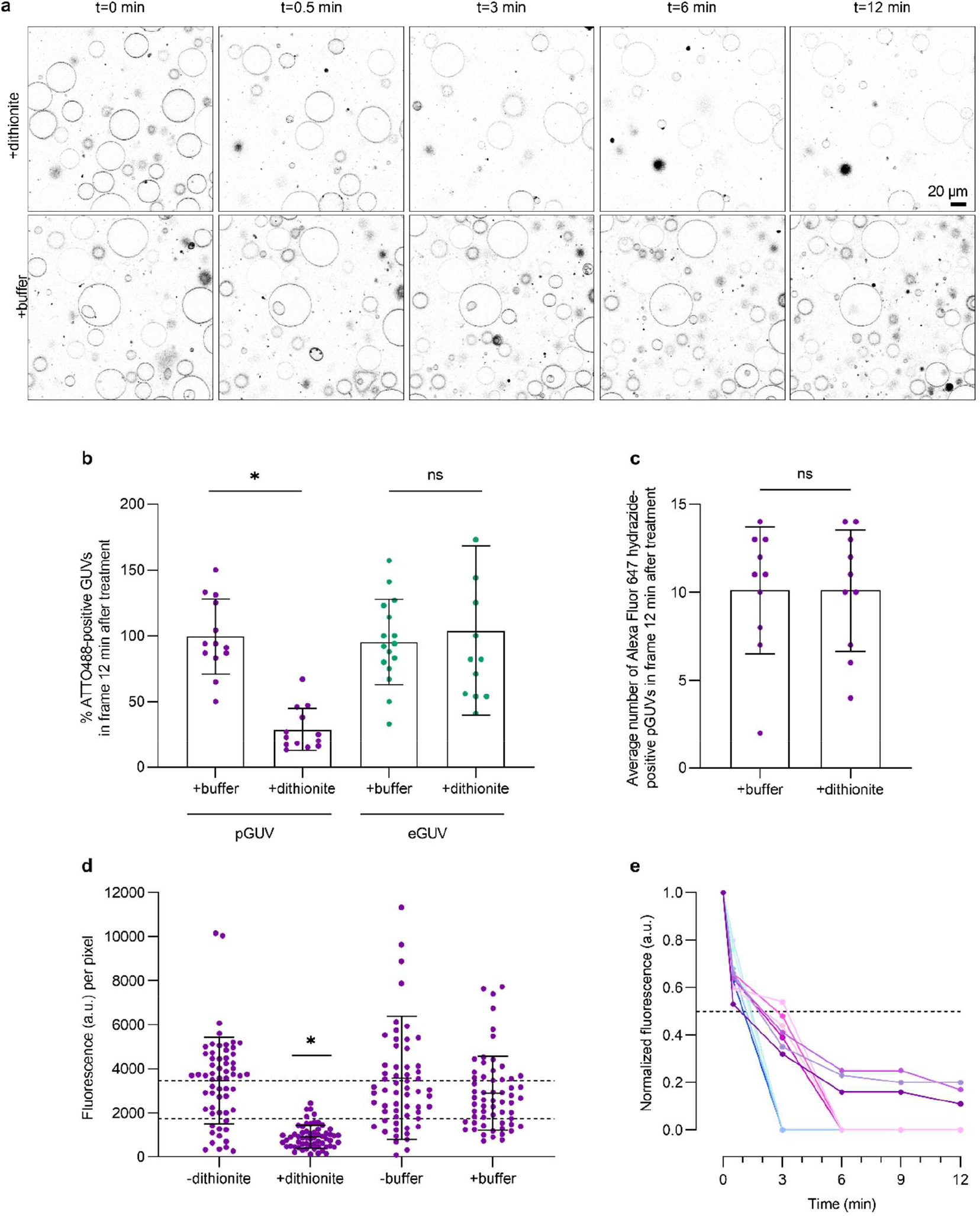
Dithionite bleaching of ATTO488-PE in pGUVs. (**a**) Confocal time series of ATTO488-PE-containing pGUVs before (t=0 min) and after (t=0.5, 3, 6, 12 min) addition of dithionite or buffer. (**b**) Dot plot showing the percentage of ATTO488-PE-positive pGUVs and eGUVs in more than ten fields of view. at t=12 min, compared with the value at t=0 min in the same field. (**c**) Dot plot showing the average number per imaged field of Alexa Fluor 647 hydrazine (lumenal marker)-positive pGUVs and eGUVs after 12 min treatment with dithionite or buffer. (**d**) Dot plot of ATTO488-PE fluorescence of individual pGUVs within the field of view before and 12 min after dithionite or buffer treatment (purple dots, n=60). Black lines indicate mean ± s.d. (one-way ANOVA and Tukey test, *\*P* < 0.0001). Dashed lines are provided as a guide to indicate 100% and 50% levels of fluorescence based on the mean value at t=0 min. (**e**) Dithionite reduction of ATTO488-PE fluorescence of eleven individually tracked pGUVs (coded in different colors), normalized to the fluorescence value at t=0 min. The arrowhead indicates dithionite addition. The horizontal dashed line corresponds to 50% loss of fluorescence. Four of the traces show complete loss of fluorescence by t=3 min, four show complete loss of fluorescence by t=6 min, and three show detectable fluorescence at t=12 min.

We tracked the rate of fluorescence loss in individual pGUVs upon addition of dithionite and observed that some pGUVs showed a very fast loss of fluorescence, within ~3 min. Others lost fluorescence more slowly, going ‘dark’ by 6 min or remaining detectable even at t=12 min. These latter vesicles had only 16 ± 4.5% (n=3) of their starting fluorescence remaining at t=12 min (Fig. 6e). Indeed, a more robust sampling of these vesicles indicated that at the 12-min time point they possessed, on average, ~26% of their starting value of ATTO488-PE fluorescence (Fig. 6d). This behavior is distinct from that observed for eGUVs or predicted for pGUVs that lack a scramblase - in these cases, the extent of fluorescence reduction would be expected to be close to 50% (see above). It is therefore likely that the pGUVs that are still detectable at t=12 min correspond to kinetic intermediates in the scrambling process. As presented in the Discussion section, slow scrambling in these vesicles is likely because they contain relatively few scramblases. We conclude that all the pGUVs in our sample are scramblase positive, with the majority demonstrating complete bleaching of the reporter lipid within 3-6 min of dithionite addition.

## Discussion

We report a method to generate pGUVs containing functional ER phospholipid scramblases and present a fluorescence-based characterization of scramblase activity at the single vesicle level using a photostable fluorescent phospholipid as transport reporter. GUVs provide an elegant experimental system for studies of lipid transporters, as their cell-like size permits observations by light microscopy techniques of individual vesicles. Realizing the potential of this system for the study of lipid transporters requires sophisticated flip-flop assays in combination with efficient protein incorporation into GUVs under conditions that preserve protein activity while still allowing a high yield of GUVs. In this study, we established a phospholipid scramblase assay for single GUV analysis based on the irreversible elimination of the fluorescence of an ATTO-dye labeled phospholipid, ATTO488-PE, with sodium dithionite. Our results show that this photostable lipid derivative readily reacts with dithionite and is transported by ER scramblases, thus providing a sensitive tool for studying scramblase activity at single vesicle level. Notably, the fast kinetics of dithionite-mediated reduction of ATTO488-PE might be helpful in the future for monitoring transbilayer distribution and movement of lipid analogs with high time resolution, for example in conjunction with stopped-flow techniques ^35^.

To reconstitute ER membrane proteins in GUVs, we used the electroformation method, where aqueous dispersions of proteoliposomes with fluorescently labeled yeast ER proteins were deposited on ITO plates followed by application of an AC-electric field. In line with previous results ^22,24,27,36–39^, this approach resulted in effective formation of thousands of pGUVs per chamber. Under these conditions, a small fraction of multilamellar and/or tubular vesicles was detected by fluorescence microscopy and could be excluded from analyses. The pGUVs were prepared from proteoliposomes of relatively simple lipid composition, although higher lipid complexity can be achieved by mixing the proteoliposomes with liposome solutions of different compositions prior to electroformation ^26^. As reconstitution of membrane proteins is not always efficient in complex lipid mixtures, especially those containing cholesterol, this procedure would allow future study of the effect of complex lipid mixtures on protein activity by reconstituting the proteins efficiently into ‘simple liposomes’ and introducing the more complex lipid environment during GUV formation. This is an objective for future work.

The cell-size diameter of the GUVs allows for single vesicle analysis. Whereas ATTO488-PE bleaching in eGUVs occurred relatively uniformly with a t_½_ of approximately 1 min reaching an expected extent of reduction close to 50%, the time course of complete fluorescence loss upon dithionite addition differed between individual pGUVs. This can be explained by differences in the number of scramblases per pGUV, with the likely rate-limiting step being scramblase-mediated lipid translocation across the pGUV bilayer. Thus, if we consider a 30 µm diameter pGUV with a single scramblase, the mean time it would take for a freely diffusing phospholipid to encounter the scramblase and be captured by it is ~45 min (see Materials and Methods). Given the number of phospholipid molecules in one leaflet of the pGUV, this yields a capture frequency of ~10^8^ min^−1^. In comparison, the unitary rate of phospholipid scramblases is reported as >10^5^ s^−1^ or roughly ~10^7^ min^−1^. Thus, a single scramblase in a 30 µm diameter pGUV would equilibrate the phospholipid populations of the two leaflets of the bilayer in several hundred minutes. In order to achieve the rapid loss of ATTO488-PE fluorescence that we report in individual pGUVs, we estimate that each pGUV must be reconstituted with ten or more scramblases. This is more than reasonable considering the number of proteoliposomes that contribute to the electroformation of a single pGUV. Future work will evaluate the kinetics of scrambling as a function of the number of reconstituted scramblases, a parameter that can be manipulated by carrying out electroformation using proteoliposomes diluted with protein-free liposomes.

In this study, GUV formation was performed under low salt conditions ^22,25^, which did not affect the activity of the ER scramblases. The electroformation technique under low salt conditions has also been shown to be effective in incorporating and maintaining the activity of the sarcoplasmic reticulum Ca^2+^-ATPase and the H^+^ pump bacteriorhodopsin in GUVs ^22^. However, activity may not be preserved for all membrane transporters under these conditions. Notably, GUVs can be effectively prepared in buffers containing physiological salt concentrations by applying a voltage with a higher frequency (~500 Hz) ^40^. Furthermore, protocols have been established for functional membrane protein reconstitution into GUVs using detergent-destabilization of preformed GUVs ^41^, spontaneous fusion of liposomes ^42^, fusion via osmotic shock ^43^, charge-mediation ^44,45^, peptide induced fusion ^46^, and gel-assisted swelling ^23^. Thus, our assay should be broadly applicable without imposing serious constraints either on GUV composition nor on buffer solution.

## Materials and Methods

### Materials

L-α-phosphatidylcholine (egg PC), 1-palmitoyl-2-{6-[(7-nitro-2-1,3-benzoxadiazol-4-yl)amino]hexanoyl}-*sn*-glycero-3-phosphocholine (C6-NBD-PC), and 1,2-dioleoyl-*sn*-glycero-3-phosphoetholamine-N-(cap biotinyl) (Biotinyl-PE) were obtained from Avanti Polar Lipids Inc. (Birmingham, AL, USA). 1,2-dioleoyl-*sn*-glycero-3-phosphoethanolamine headgroup labeled with ATTO 488 (ATTO488-PE) was obtained from ATTO-TEC GmbH (Siegen, Germany). Bio-Beads SM-2 Resin and Bio-Gel P-6 were obtained from Bio-Rad Laboratories Inc. (Hercules, CA, USA). Unless indicated otherwise, all other chemicals and reagents were obtained from Sigma-Aldrich (München, Germany). Protease inhibitor cocktail contained aprotinin (5 mg), leupeptin (5 mg), pepstatin (5 mg), antipain (25 mg) and benzamidine (785 mg) in 5 ml DMSO used at 1:1000 dilution. All buffers and solutions used for vesicles were filter-sterilized through a polyethersulfone membrane with a pore size of 0.2 µm (Filtropur, Sarstedt AG & Co. KG, Nümbrecht, Germany).

### Preparation of detergent-solubilized ER membrane proteins (‘Triton Extract’, TE)

TE was prepared from yeast cells (BY4741 strain) as previously described ^9,47^, except for an additional salt-wash step of the membranes prior to detergent solubilization. Briefly, cells harvested at OD_600_ ~2 were washed and homogenized using glass beads (0.5 mm) in ice-cold lysis buffer (10 mM HEPES-NaOH pH 7.4, 10 mM MgCl_2_, 5% (w/v) glycerol, 1 mM dithiothreitol) supplemented with protease inhibitor cocktail. After a low-speed centrifugation (2,500 × *g*_*av*_, 10 min, 4° C), the supernatant was centrifuged at 200,000 × *g*_*av*_ (30 min, 4° C) to pellet membranes. The membranes were resuspended in ice-cold resuspension buffer (10 mM HEPES-NaOH pH 7.4, 100 mM NaCl, protease inhibitor mixture) containing 0.5 M sodium acetate and incubated on ice for 30 min. Membranes were re-pelleted, resuspended in ice-cold resuspension buffer and solubilized by gradual addition of an equal volume. of ice-cold extraction buffer (10 mM HEPES-NaOH pH 7.4, 100 mM NaCl, 2% (w/v) Triton X-100, 0.5 mM phenylmethylsulfonyl fluoride, protease inhibitor cocktail) with a final concentration of 1% (w/v) Triton X-100. Samples were incubated on ice for 30 min before insoluble material was removed by centrifugation (200,000 × *g*_*av*_, 1 h, 4° C) to generate a clear supernatant (TE), which was collected, aliquoted, snap frozen in liquid nitrogen and stored at −80° C. Protein content was determined by the micro bicinchoninic acid protein assay kit (Thermo Fisher Scientific, Rockford, IL, USA).

### Labeling TE proteins with a fluorescent dye

TE proteins were fluorescently labeled with the amine-reactive dye Alexa Fluor 647 NHS-Ester (Thermo Fisher Scientific, Rockford, IL, USA) as follows. TE (95 µg) was mixed with 5.7 nmol of Alexa Fluor 647 NHS-Ester in labeling buffer (2 mM HEPES-NaOH pH 8.3, 1 mM NaCl, 1% (w/v) Triton X-100) for 1 h under end-over-end mixing. Unreacted dye was removed by two passages over an equilibrated Bio-spin® 6 column filled with Bio-gel P-6 media in labeling buffer and centrifuged in a tabletop centrifuge (Eppendorf 5810 R, rotor A-4-62) at 1500 rpm for 3 min. The eluate (termed TE647) was used immediately for proteoliposome reconstitution.

### Reconstitution of proteoliposomes

Proteoliposomes were prepared from a mixture of unlabeled TE or TE647 and Triton X-100-solubilized phospholipids as previously described ^9^. Briefly, egg PC (4.5 µmol), fluorescent lipids (0.1 mol% ATTO488-PE or 0.3 mol% C6-NBD-PC), and biotinyl-PE (1 mol%) in chloroform were dried under nitrogen in a glass screw-cap tube and then dissolved in Triton X-100 containing buffer (2 mM HEPES-NaOH pH 7.4, 1 mM NaCl, 1% (w/v) Triton X-100). TE was added to solubilized lipids to a protein/phospholipid ratio of 22 mg mmol^−1^. Protein-free liposomes were prepared similarly by replacing TE with buffer. Liposome formation was induced by detergent removal using Bio-Beads SM-2 (prewashed with methanol, water, and buffer) over two stages: 100 mg Bio-Beads SM-2 incubation with end-over-end mixing for 3 h followed by additional 200 mg Bio-Beads SM-2 with end-over-end mixing overnight at 4° C. The resulting vesicles were collected and stored at 4° C until use.

### Liposome analysis

Protein reconstitution was detected by SDS-PAGE under reducing conditions on a 10% polyacrylamide gel, followed by in-gel fluorescence scanning of Alexa 647 (ChemiDoc™ MP device, Bio-Rad Laboratories GmbH, München, Germany) and silver staining (GE Healthcare, Uppsala, Sweden). The phospholipid recovery in reconstituted liposomes was 70-90%, as described previously ^9^. More than 99.98% of the initial Triton X-100 amount was removed, as determined by extraction with four volumes of chloroform/methanol (1/2, v/v) and measurement of the absorbance of the supernatant at 275 nm ^48^. To assay the flip-flop of fluorescent lipids, the fluorescence of the vesicles was measured at 25° C with a fluorometer equipped with magnetic stirring (PTI-Quantamaster 800, Horiba, Benzheim, Germany). Proteoliposomes (50 µl) were diluted into 2 ml (final) of external solution containing 2 mM HEPES-NaOH pH 7.4, 1 mM NaCl. Fluorescence traces were recorded for ATTO488 at λ_ex_/λ_em_ 507/530 nm, and for NBD at λ_ex_/λ_em_ 470/534 nm, for 300 s, slit width 4-5 nm, resolution 0.1 s. Sodium dithionite (Thermo Fisher Scientific) was added after 30 s to the cuvette to start the reaction (40 µl of a 1 M stock solution prepared in 0.5 M unbuffered Tris; 20 mM final concentration). Data were collected using FelixGx 4.9.0 at a sampling rate of 1 Hz. To confirm complete reduction of all analogues by dithionite, Triton X-100 was added to a final concentration of 0.5% (w/v). The percentage of fluorescent lipid that was reduced (F_red_) was calculated as (F − F_end_)/(F_start_ − F_end_) × 100, where F is the fluorescence of liposomes at the given time, F_end_ is the last 20 s of fluorescence after Triton X-100 addition, F_start_ is initial fluorescence of liposomes 15 s before dithionite addition. The half-times for dithionite reduction were obtained by nonlinear regression curve fitting to a one-phase exponential decay curve using Prism version 9.1.0 (GraphPad Software, San Diego, CA).

### Preparation of giant unilamellar vesicles (GUVs)

GUVs were generated from proteoliposomes or liposomes in a chamber made of indium-tin-oxide-coated (ITO) glass slides (Präzisions Glas & Optik GmbH, Iserlohn, Germany) by electroformation ^22,25^. Briefly, proteoliposomes or liposomes were diluted with distilled water to a lipid concentration of 0.8 mg ml^−1^, and 50 µl of this suspension was applied as droplets on each UV-cleaned glass slide and dehydrated overnight in a sealed chamber containing a saturated NaCl solution. Subsequently, a chamber was formed by separating the ITO glass slides with a Teflon spacer containing copper electrodes. The chamber was sealed and filled with 1 ml 250 mM sucrose solution. GUVs were electroformed by applying a sinusoidal voltage for 4 h (12 or 20 Hz, 0.2-1.3 V, increasing every 6 min). Detachment of the GUVs from the slides occurred by applying a second sinusoidal voltage for 30 min (2.0 V, 4 Hz). GUVs were gently transferred and stored in a 1.5 ml tube.

### PEGylation of cover glass slides

Glass coverslips (26 × 76 mm, #1.5, Thermo Fisher Scientific) were cleaned in glass jars as described previously ^49^, except for the Piranha etching, followed by amino-silanization using 3-aminopropylthriethoxysilane. Surface PEGylation was performed according to Lamichhane et al. ^50^ using mPEG-SC (5,000 Da, Laysan Bio Inc., Arab, AL) and biotin-PEG-SC (5,000 Da, Laysan Bio Inc.), introducing 5% biotinylated PEG, and stored at −20° C until use.

### GUV immobilization

GUVs were immobilized on biotin-PEGylated cover glass slides via avidin binding. A chamber was formed by gluing a sticky-Slide 8 Well μ-Slide from Ibidi (Munich, Germany) onto the PEGylated cover glass slide using Norland Optical Adhesive 60 (Norland products, Cranbury, USA) followed by 2 min UV radiation (Ultra-Violet/Ozone Probe and Surface Decontamination Unit, novascan, Boone, IA). Avidin (200 µl, 0.025 mg ml^−1^ in distilled water) was added to each chamber. After 20 min incubation, free avidin was removed by washing four times with 250 mM glucose solution. GUVs (100 µl) were added using a cut pipet tip and the chamber was filled up to 500 µl with 250 mM glucose solution.

### Microscopy

Fluorescence microscopy and image acquisition were carried out using a Leica TCS SP8 confocal laser scanning microscope (Leitz, Wetzlar, Germany) equipped with 63x/1.20 water objective. Images were acquired using a 400 Hz unidirectional scanner, a pixel size of 246.27×246.27 μm, a pinhole of 100 μm (1 AU) with Leica HyD detectors. The λ_ex_/λ_em_ used for imaging were as follows: ATTO488 507/517-550 nm, Alexa647 645/655-780 nm, and Alexa Fluor 647 hydrazide 651/661-780 nm. Images were scanned using the same conditions of the pinhole, gain, laser power (10% for ATTO488-PE), and detector offset in each experiment.

### GUV image analysis

To assay the flip-flop of ATTO488-PE, up to thirteen images were obtained before and after dithionite treatment. Dithionite was dissolved immediately before use in 250 mM glucose solution and added as 5 µl (2 mM final concentration) to the GUV containing glass slide chamber. A time series was started immediately, obtaining five consecutive images at 0.5, 3, 6, 9 and 12 minutes. Buffer treatments were performed as described for dithionite addition. To assess dithionite permeation, GUVs were generated in 250 mM sucrose solution containing 10 µM Alexa Fluor 647 hydrazide, a soluble dye that cannot cross membranes (Thermo Fisher Scientific). Subsequently, GUVs were mixed 1:1 (v/v) in 250 mM glucose solution in a 1.5 ml tube to a final volume of 200 µl. Dithionite (100 µl) was added to a final concentration of 10 mM and the tube was gently inverted three times. GUVs were transferred to PEGylated cover glass slides using a cut pipet tip and allowed to sediment. Images was obtained 12 min after dithionite addition. The dithionite reactivity of free 10 µM Alexa Fluor 647 hydrazide (λ_ex_/λ_em_ 649/666 nm) in 250 mM sucrose solution was assayed at 25° C in a stirred cuvette using PTI-Quantamaster 800.

### Data analysis

Quantitative data are presented as mean ± standard deviation (s.d.). The data was analyzed with ANOVA followed by Šídák’s multiple-comparison test or two-tailed Student *t* test. The statistical analyses were carried out using GraphPad Prism version 9.1.0. The number of replicates is reported in figure captions. P values of less than 0.05 were regarded as statistically significant.

### Calculations

We estimated the mean time (t_dc_ or ‘mean time to capture’) it would take for a phospholipid diffusing in the GUV membrane to encounter a scramblase, considered here as a fixed absorbing region of radius *s*. We assume that the GUV is 30 μm in diameter and has a single scramblase, and that phospholipids diffuse in the GUV membrane with a diffusion coefficient *D*. Included in the calculation is the distance *b*, equal to half the circumference of the GUV, i.e., the distance from the scramblase at which as many lipids cross a boundary moving towards the scramblase as away from it. For the case here where *s*/*b* << 1, t_dc_ =(*b*^2^/2D)•(ln(*b*/*s*) −3/4) ^37,38^. Taking *b* = 4.71 × 10^−5^ m, *D* = 4 × 10^−12^ m^2^ s^−1^ (from Ref. 39), and *s* = 1.8 × 10^−9^ m (the cross-sectional radius of a cylindrical opsin molecule) ^40^, t_dc_ = 2610 s or 43.5 min. As the number of phospholipids (*P*) in a single leaflet of a 30 μ GUV is ~4.35 × 10^9^ (calculated from the surface area of the GUV, and assuming that the cross-sectional area of a single phospholipid is 0.65 × 10^−18^ m^2^ (from Ref. 45), the frequency of phospholipid capture by the scramblase is *P*/t_dc_ = 10^8^ min^−1^.

## Online supplemental material

Suppl. Fig. S1 outlines and validates liposome reconstitution with fluorescently labeled endoplasmic reticulum (ER) membrane proteins. Suppl. Fig. S2 provides additional data on eGUVs to support the results shown in Fig. 4b.

## Acknowledgments

We gratefully acknowledge Alice Verchére and Anne-Mette Petersen for assistance in TE purification, Eckhard Hofmann and Christian Herrmann (Ruhr University, Bochum) for access to instrumentation, Robert Tampé and Tim Diederichs (Goethe University, Frankfurt) for helpful discussions and technical advice on the project, and André Nadler (MPI-CBG Dresden) for discussion. This work was supported by DAAD Grant 57386621 (to T.G.P.) and the German Research Foundation (GU 1133/11-1; INST 213/886-1 FUGG to T.G.P).

## Author contributions

P.P.M.M., A.K.M., and T.G.P. conceived and designed the project. T.G.P. and A.K.M supervised the research. P.P.M.M. performed and analyzed the experiments. P.P.M.M. wrote the first version of the manuscript and revised it with the help of A.K.M. and T.G.P. All authors reviewed the final version.

---

**Supplementary figure S1.**
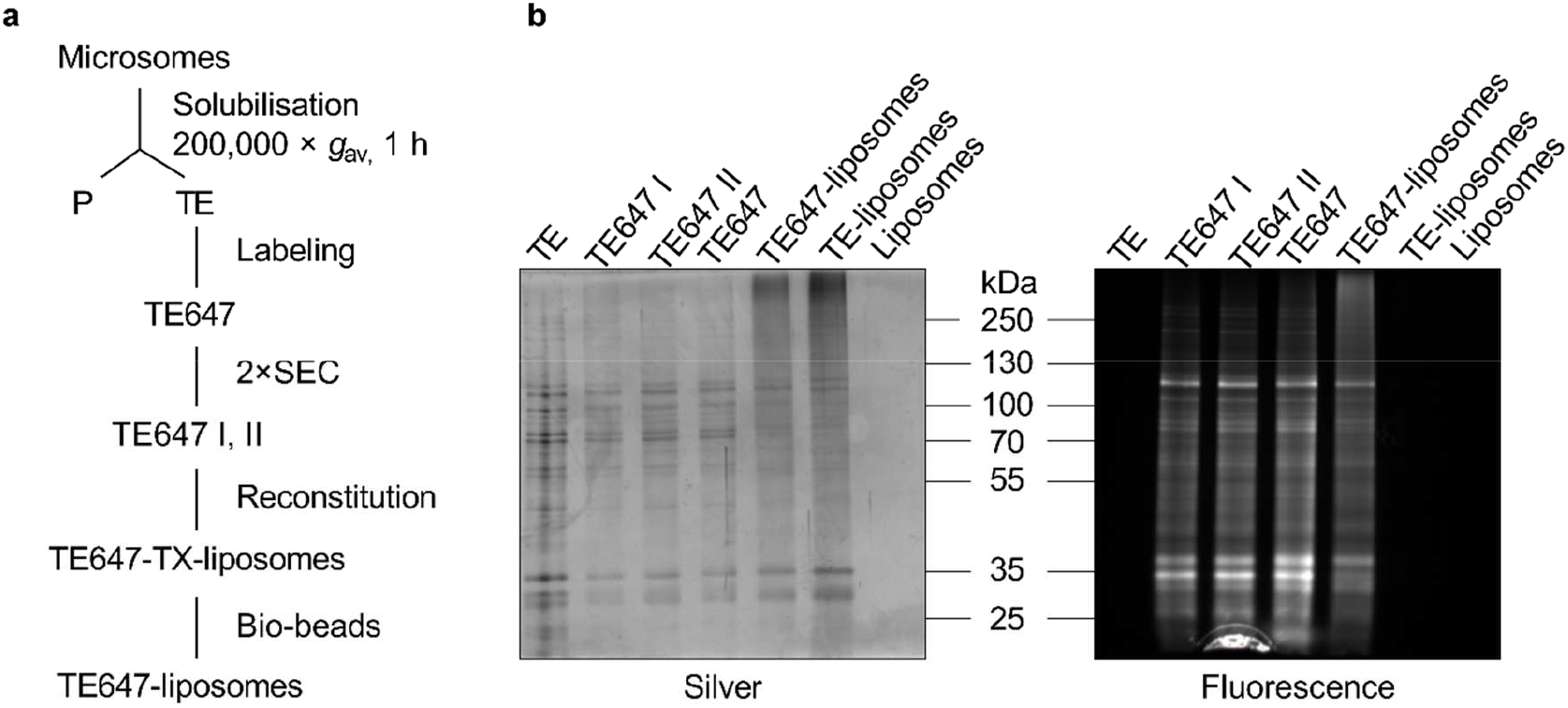
Preparation of fluorescently labeled endoplasmic reticulum (ER) membrane proteins and reconstitution into large unilamellar liposomes. (**a**) Experimental workflow. ER membrane proteins were selectively solubilized from yeast microsomes with Triton X-100 to generate a ‘Triton Extract’ (TE). TE was fluorescently labeled (TE647) with Alexa Fluor 647 NHS ester, excess fluorophore was removed via two rounds of size-exclusion chromatography (SEC) (yielding TE647 I and subsequently TE647 II), and the resulting labeled proteins were reconstituted together with egg PC, ATTO488-PE, and Biotinyl-PE into liposomes (TX-liposomes) using Bio-beads SM2 to remove detergent. Proteoliposomes were also generated with unlabeled TE and protein-free liposomes were generated in parallel. (**b**) SDS-PAGE analysis of samples from different steps during proteoliposome reconstitution. The gel was visualized by silver staining (left) and fluorescence scan of Alexa Fluor 647 (right).

**Supplementary figure S2.**
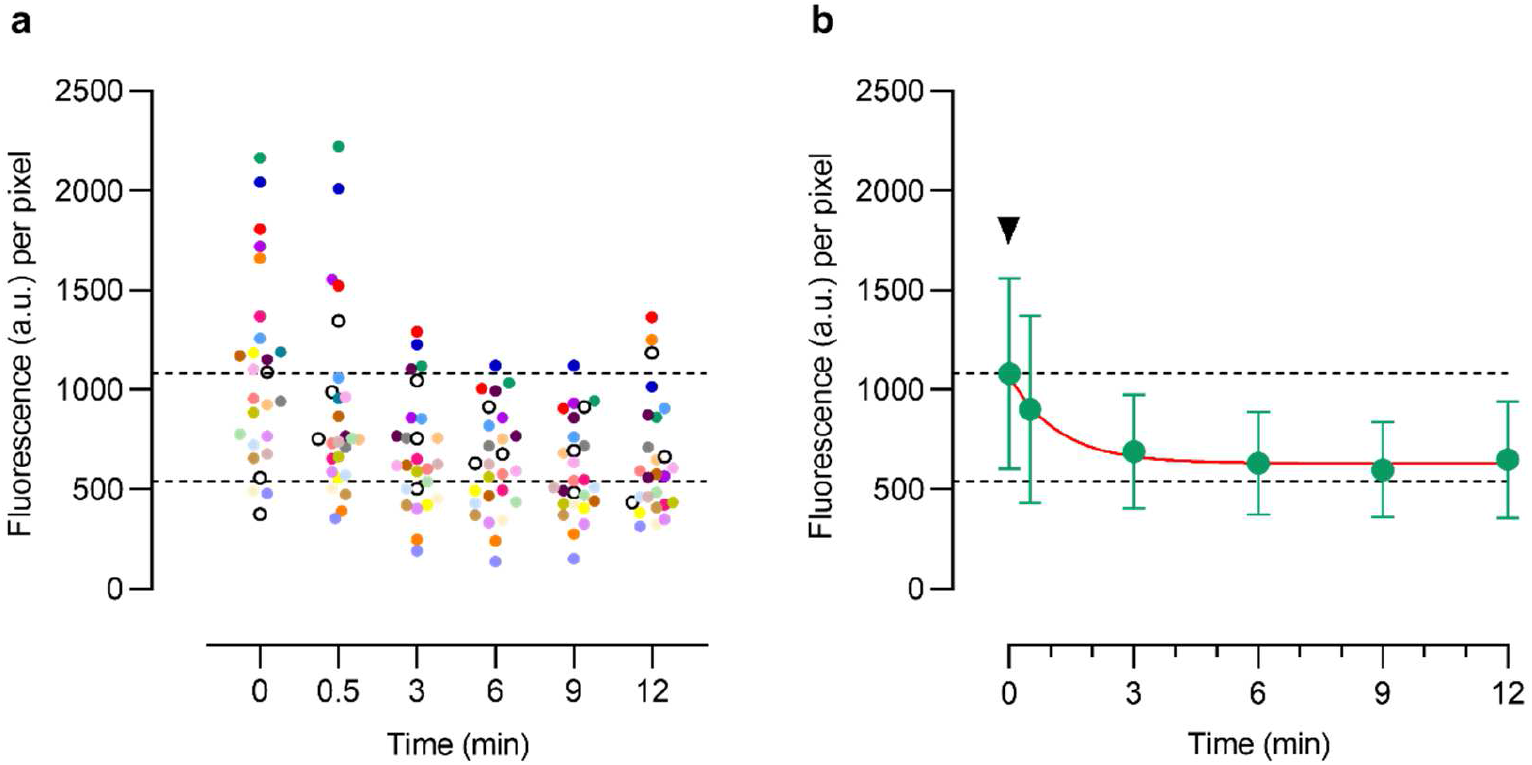
Dithionite bleaching of ATTO488-PE in empty GUVs. (**a**) Dot plot of ATTO488-PE fluorescence intensities of individual eGUVs (each eGUV is uniquely color coded, n=26) before (t=0 min) and after dithionite addition (t=0.5, 3, 6, 9 and 12 min). Black open circles indicate three eGUVs that did not react to dithionite. Dashed lines indicate 100% and 50% levels of fluorescence based on the mean value at t=0. (**b**) Time course of the loss of average ATTO488-PE fluorescence intensity of eGUVs upon addition of dithionite (arrowhead). Data are compiled from panel a and presented as mean ± s.d. (n=26). The red line represents a monoexponential fit of the data (t_½_ = 0.82 min). Dashed lines indicate 100% and 50% levels of fluorescence based on the mean value at t=0 min.

